# Septins regulate border cell shape and surface geometry downstream of Rho

**DOI:** 10.1101/2021.04.08.439079

**Authors:** Allison M. Gabbert, James A. Mondo, Joseph P. Campanale, Noah P. Mitchell, Adele Myers, Sebastian J. Streichan, Nina Miolane, Denise J. Montell

**Affiliations:** Molecular, Cellular, and Developmental Biology Department at University of California, Santa Barbara; Kavli Institute for Theoretical Physics at University of California, Santa Barbara; Physics Department at University of California, Santa Barbara; Electrical and Computer Engineering Department at University of California, Santa Barbara

## Abstract

Septins self-assemble into polymers that bind and deform membranes *in vitro* and regulate diverse cell behaviors *in vivo*. How their *in vitro* properties relate to their *in vivo* functions is under active investigation. Here we uncover requirements for septins in detachment and motility of border cell clusters in the *Drosophila* ovary. Septins and myosin colocalize dynamically at the cluster periphery and share phenotypes, but surprisingly do not impact each other. Instead, Rho independently regulates myosin activity and septin localization. Active Rho recruits septins to membranes while inactive Rho sequesters septins in the cytoplasm. Mathematical analyses reveal how manipulating septin expression alters cluster shape and surface geometry. This study shows that the level of septin expression regulates surface properties at different scales. This work suggests that downstream of Rho, septins tune surface deformability while myosin controls contractility, the combination of which govern cluster shape and movement.

## Introduction

Cell migration is essential for development, wound healing, immune responses, and tumor metastasis [reviewed in (Perez-Vale and Peifer, 2020; Ridley et al., 2003; SenGupta et al., 2021; Stuelten et al., 2018; Yamada and Sixt, 2019)]. While our fundamental understanding of the molecular mechanisms controlling cell motility derives primarily from studying cells migrating individually on glass, *in vivo*, cells frequently move collectively [reviewed in (Haeger et al., 2015; Mishra et al., 2019a; Scarpa and Mayor, 2016; Shellard and Mayor, 2021)]. *In vivo*, cells also move through complex, cell-rich microenvironments that are difficult if not impossible to recapitulate *in vitro*. Therefore, *in vivo* models amenable to genetic analysis and live imaging are important.

Border cells in the *Drosophila* egg chamber provide an excellent model to study collective cell migration (Friedl and Gilmour, 2009; Montell, 2003; Montell et al., 2012; Prasad et al., 2011; Rørth, 2002). The border cell cluster is composed of four to six migratory cells that surround and transport two non-motile polar cells from the anterior end of the egg chamber in between nurse cells, to the oocyte during oogenesis [reviewed in (Montell et al., 2012)]. Cytoskeletal dynamics are critical determinants of cell shape and movement in general (Seetharaman and Etienne-Manneville, 2020; Zegers and Friedl, 2014) and border cells in particular (Chang et al., 2018; Combedazou et al., 2017; Mishra et al., 2019b; Plutoni et al., 2019; Sawant et al., 2018; Wang et al., 2020; Zeledon et al., 2019), where the *in vivo* requirement for the small GTPase Rac in F-actin-rich protrusion and migration was first demonstrated (Murphy and Montell, 1996).

F-actin, microtubules, and intermediate filaments are well-studied, dynamic polymers that contribute to cell shape and motility (Seetharaman and Etienne-Manneville, 2020). Septins are filament-forming GTPases that have been described as a fourth major cytoskeletal element (Mostowy and Cossart, 2012). First discovered as a key component required for septation in budding yeast (Hartwell, 1971), septins are now known to be conserved throughout animals and fungi where they commonly localize to the cell cortex and act as protein scaffolds (Woods and Gladfelter, 2021). Septins regulate cell polarity, ciliogenesis, cytokinesis, individual and collective cell migrations, and other processes (Tooley et al., 2009; Shinoda et al., 2010; Oh and Bi, 2011; Olguin-Olguin et al., 2021; Shindo and Wallingford, 2014). In addition to their functions in normal cellular processes, alterations in septin expression have been implicated in diseases including neurodegenerative diseases such as Alzheimer’s and Parkinson’s, many types of cancer such as mixed lineage leukemia and ovarian cancer, and male infertility (Dolat et al., 2014; Hall and Russell, 2004; Marttinen et al., 2015; Peterson and Petty, 2010; Saarikangas and Barral, 2011), though the precise contributions of septins to disease etiology are not yet clear. Key open questions in the septin field include how the *in vitro* properties of septins relate to their *in vivo* functions, how alterations in septin expression levels affect their functions, and how septin activity is regulated *in vivo*?

Fruit flies have a simplified set of five septins compared to 13 encoded in the human genome. *Drosophila* Sep1, Sep2, and Pnut have been purified as a complex (Field et al., 1996). In an RNA expression screen, we previously reported that mRNAs encoding Sep1, Sep2, and Pnut are expressed and/or enriched in border cells (Wang et al., 2006). Here we show that Sep1, Sep2 and Pnut proteins are expressed in both cytoplasmic and membrane associated pools, and that the active form of the Rho1 GTPase recruits septins to border cell membranes whereas dominant-negative Rho sequesters septins in the cytoplasm. We report that septins are required for border cell migration and that their knockdown smoothes border cell surfaces, whereas their overexpression roughens them. Knockdown and overexpression also change the overall shape of the cluster and impair motility. Together these results demonstrate that septin levels tune cell surface topography and, when expressed at precisely appropriate levels, promote collective motility downstream of Rho.

## Results

### Septins colocalize in collectively migrating border cells

Individual septin monomers contain a proline-rich amino-terminal region, a central core that includes a GTP-binding domain, and a carboxy-terminal region (Figure 1A). Septins oligomerize via interactions between the GTP-binding domains (G-G) and through the interactions between N- and C-termini of different subunits (N-C) (Figure 1B). Septin polymers can form filaments, bundles, and more complex structures such as gauzes, bundles, and rings (Figure 1C) (Bridges et al., 2016). *Drosophila* and human septins show between 55 and 73% amino acid sequence identity (Figure S1A).

**Figure 1:**
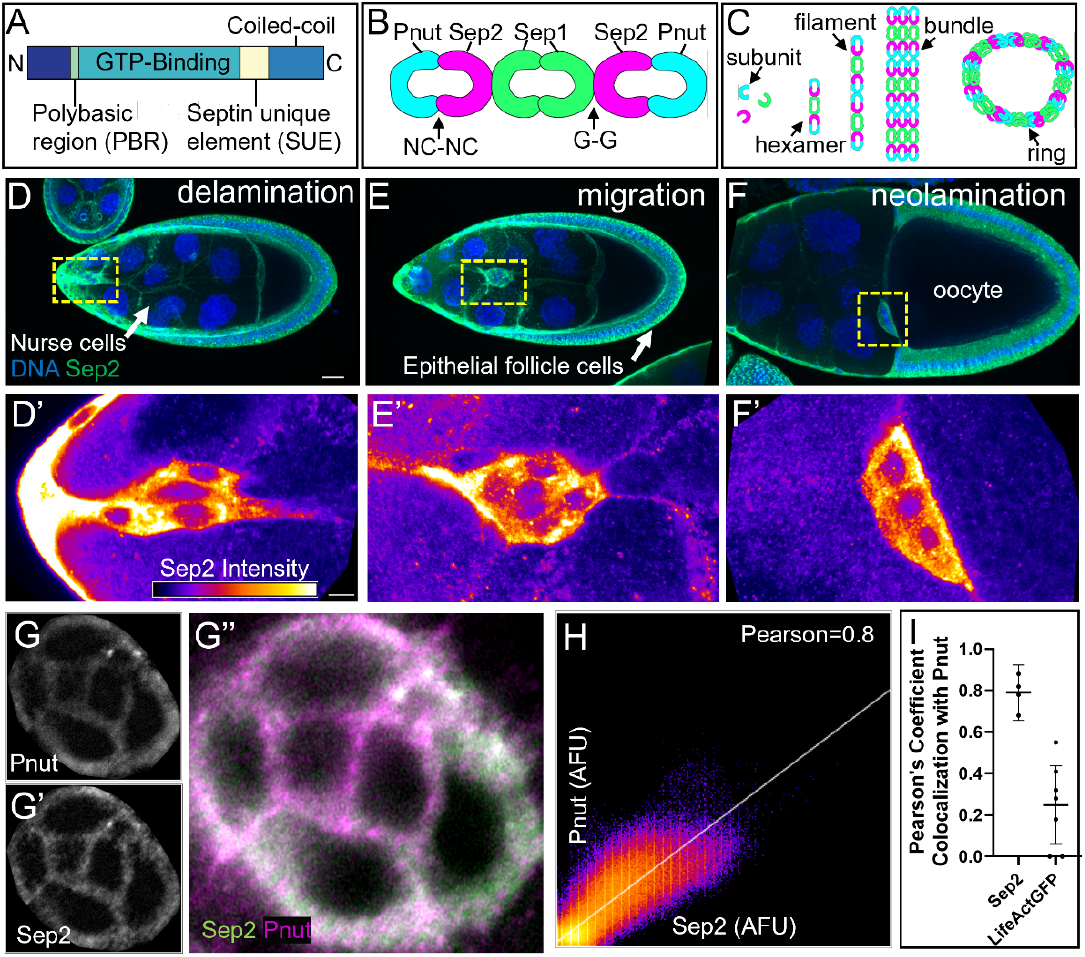
Septins colocalize in collectively migrating border cells. (A) Schematic of septin domains. (B) Septins assemble into complexes through interactions between their N-C and G-G interfaces. In *Drosophila*, Pnut, Sep2, and Sep1 form a complex as shown. (C) Septin subunits can form higher order structures. (D-F) Max intensity projections of egg chambers labeled with Hoechst (blue) and Sep2:GFP (green). (D’-F’) Single slices of the border cell clusters from D-F with Sep2:GFP shown in fire LUT. (G-G’’) Single slice of a border cell cluster imaged by Airyscan labeled with Pnut (G), Sep2 (G’), or both (G’’). (H-I) Graph of mean Pnut and Sep2 intensities showing co-localization. Error bars are 95% confidence intervals.. Each dot represents a single cluster.. Scale bar in D is 20 μm and 5 μm in D’.

*Drosophila* egg chambers are composed of ∼850 somatic follicle cells that surround 16 germline cells – 15 polyploid nurse cells and one oocyte (Figures 1D-1F). Border cells develop at the anterior end of the egg chamber and during stage 9 extend protrusions between the nurse cells, delaminate from the epithelium, and migrate down the central path until they reach the oocyte at stage 10 (Figures 1D-1F). To probe the expression and localization of septins in egg chambers, we examined a transgenic insertion of Sep2, expressed under its own regulatory sequences, and tagged with GFP. Sep2::GFP is expressed in all follicle cells, including border cells, throughout their migration (Figures 1D-1F and 1D’-1F’). To compare the Sep2::GFP and Pnut expression patterns, we stained egg chambers expressing Sep2::GFP for Pnut and used Airyscan confocal imaging (Figures 1G-1G’’). Sep2 and Pnut significantly colocalized with each other (Figures 1H-1I), consistent with previous reports of Sep2 assembly with Sep1 and Pnut into filaments (Fares et al., 1995; Field et al., 1996; Huijbregts et al., 2009).

### Septins are required for border cell migration

To test if septins play a role in border cell migration, we knocked down Sep1, Sep2 or Pnut using UAS-RNAi constructs. RNAi hairpins were expressed in combination with a UAS-LifeActGFP marker using c306Gal4. In egg chambers expressing the control (white) RNAi, the border cell cluster completes migration to the oocyte by stage 10 (Figure 2A). Expressing RNAi against Sep1, Sep2, or Pnut impaired migration (Figures 2B-2D). Interestingly, overexpression of Sep1 or Sep2 using c306Gal4 also disrupted migration (Figures 2E-2F). We quantified the phenotypes using categories that denote how far the cluster migrated (Figures 2G-2H). Thus, Septins 1, 2 and Pnut are required for border cell migration and the level of septin expression must be regulated, with too much or too little septin impeding motility.

**Figure 2:**
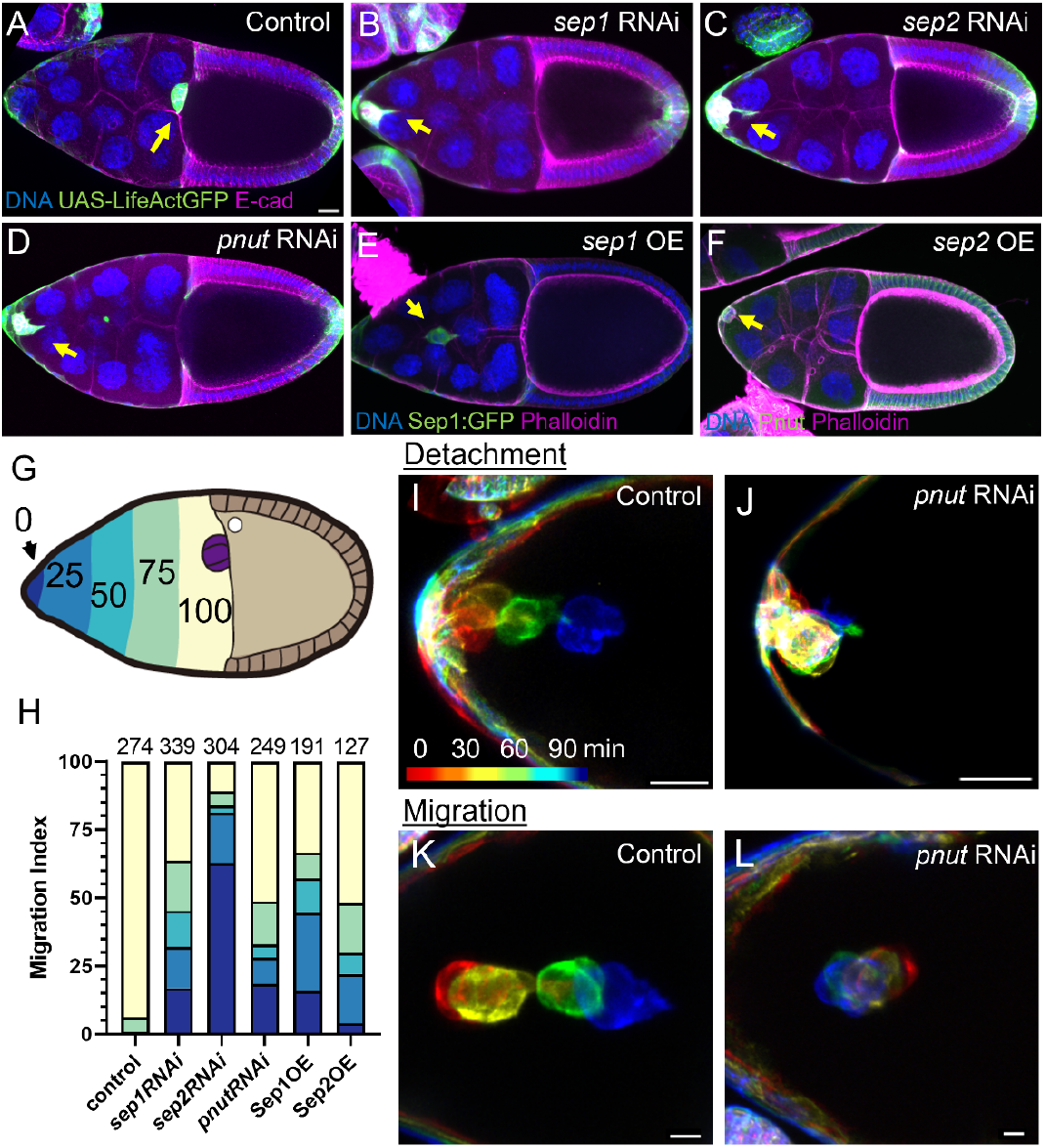
Septins are required for border cell migration. (A-D) Max intensity projections of stage 10 egg chambers expressing UAS-whiteRNAi (control) or septinRNAi and UAS-LifeActGFP in the border cell cluster by c306Gal4. Hoechst (blue) and E-cadherin (magenta). (E-F) Projections of stage 10 egg chambers expressing UAS-Sep1:GFP (E) and UAS-Sep2 (F) by c306Gal4. Hoechst (blue) and Phalloidin (magenta). Arrows (A-F) denote border cell cluster. (G) Schematic of a stage 10 egg chamber. Colors indicate percentage of the migration path. (H) Quantification of border cell migration in egg chambers after RNAi knockdown or overexpression of the indicated septins. The control was whiteRNAi. (I-L) Temporally color coded projections from movies of border cell detachment (I-J) or migration (K-L). Control (whiteRNAi) or septin knockdown clusters. Scale bars (A, I-L): 20 μm.

Border cell migration is a multistep process that begins with their specification and rounding up, followed by a delamination phase, in which the cluster detaches from the anterior epithelium (Figure 2I and Movie S1). Upon Sep1, 2, or Pnut knockdown, the border cell cluster still rounded up and the cells upregulated expression of the marker Singed, which is the fly homolog of the actin binding protein Fascin (Figures S2A-S2D). However, septin knockdown impaired detachment (Figure 2J, Movie S2). Septin overexpression also caused detachment defects (Figure S2E and Movie S3). Delamination is normally followed by migration, in which the cluster actively squeezes in between the nurse cells towards the oocyte (Figure 2K and Movie S4) (Prasad and Montell, 2007). Of those septin knockdown or overexpression clusters that detached, many (30-70%) failed to migrate normally (Figure 2L, S2F, Movie S5, Movie S6). Therefore, optimal septin expression is required for detachment and migration.

### Septins 1, 2, and Pnut are interdependent

Septins assemble into heteromeric complexes in many contexts (Dolat et al., 2014; Woods and Gladfelter, 2021), and loss of a single septin can lead to destabilization of the complex (Akhmetova et al., 2018; Menon et al., 2014; Xu et al., 2018). To test if the septin protein abundances are dependent on one another in follicle cells, we expressed Sep1, Sep2 or Pnut RNAi in clones within the follicular epithelium and stained with an antibody against Pnut. Sep1 and Sep2 reduced Pnut protein levels compared to neighboring control cells (Figures 3A-3D’, 3I). We observed similar results within the border cell cluster (Figures 3E-3H’, 3J). Quantification of the results showed that while Pnut knockdown reduced Pnut expression the most, knockdown of either Sep1 or Sep2 significantly reduced Pnut expression (Figure 3I and 3J). Similarly, border cells homozygous mutant for Sep2 exhibited reduced levels of Pnut in clonal cells (Figures S3A-S3B’). These results establish the effectiveness of the RNAi knockdowns and suggest that Sep1, Sep2, and Pnut are interdependent.

**Figure 3:**
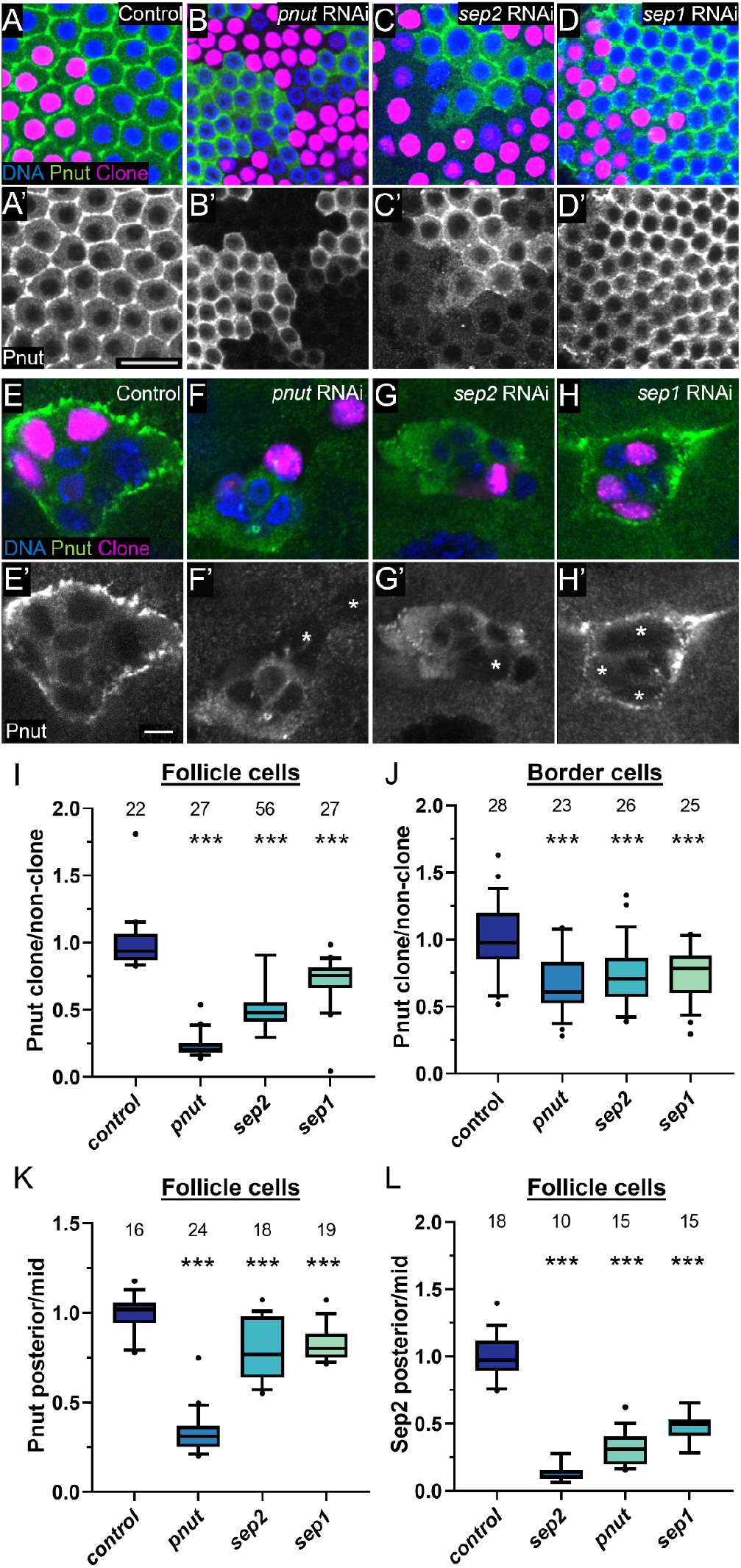
Septins 1, 2, and Pnut are interdependent. (A-D) Images of basal surface of epithelial follicle cells or (E-H) border cells clonally expressing UAS-whiteRNAi (control) or septinRNAi labeled with Hoechst (blue) and Pnut (green). Clones expressing RNAi and UAS-nuclear-localized RedStinger (magenta). (A’-D’) and (E’-H’) Pnut staining (gray) from (A-D) and (E-H) respectively. Border cell clones are marked with asterisks. (I-L) Box plot of Pnut intensity in clones compared to non-clones for (I) Follicle cells, (J) border cells, and (K) posterior follicle cells (L) Quantification of Sep2:GFP intensity in posterior follicle cells expressing RNAi compared to follicle cells not expressing RNAi. (I-L) Three asterisks represent P<0.001 when analyzed by an Ordinary one-way ANOVA, followed by Tukey post-hoc analysis. Scale bars: 20μm (A’) and 5μm (E’).

Similarly, c306Gal4-driven septin-RNAi against Sep1 or Sep2 in the posterior follicle cells significantly decreased Pnut expression (Figure 3K and Figures S3C-S3F). Additionally, knockdown of Sep1 and Pnut using c306Gal4 in a Sep2::GFP background significantly decreased GFP expression in the posterior follicle cells (Figure 3L and Figures S3G-S3J). Therefore, we conclude that Sep1, Sep2, and Pnut depend on each other, and are likely to function in a complex.

### Septins and myosin colocalize dynamically

At the yeast bud neck, active Cdc42 recruits septins (Caviston et al., 2003), which in turn recruit myosin (Schneider et al., 2013). To investigate the regulation of septins in border cells, we generated clones of cells expressing constitutively active or dominant negative forms of Cdc42 or one of two different Cdc42RNAi constructs in the follicular epithelium of the egg chamber (Figures S4A-S4D’) and the border cells (Figures S4E-S4H’). We did not observe any change in Pnut expression or localization in clones, compared to neighboring control cells. In addition, Cdc42 knockdown affects cluster cohesion, which is distinct from the septin loss of function phenotype in the border cells. Therefore Cdc42 and septins appear to function separately in border cells.

Myosin II was another candidate for interacting with septins, as septins scaffold myosin in yeast and mammalian cells (Joo et al., 2007; Schneider et al., 2013), Septin 7 is essential for planar polarization of active myosin during convergent extension movements in *Xenopus* gastrulation (Shindo and Wallingford, 2014), and myosin regulates border cell detachment and protrusion dynamics (Majumder et al., 2012). To observe myosin expression and localization, we used flies expressing a tagged third copy of spaghetti squash (Sqh), which encodes the regulatory light chain of nonmuscle myosin II. Myosin is highly dynamic in the migrating border cell cluster, appearing as moving “myosin flashes” (Aranjuez et al., 2016; Mishra et al., 2019b), so we conducted live imaging of clusters expressing both Sqh:mCherry (Figures 4A-4F) and Sep2::GFP (Figures 4A’-4F’). We imaged at 15 second intervals to capture the rapid myosin flashes along the periphery. Sep2 and Sqh traveled together dynamically (Figure 4G). This dynamic colocalization suggested a possible link between septins and myosin.

**Figure 4:**
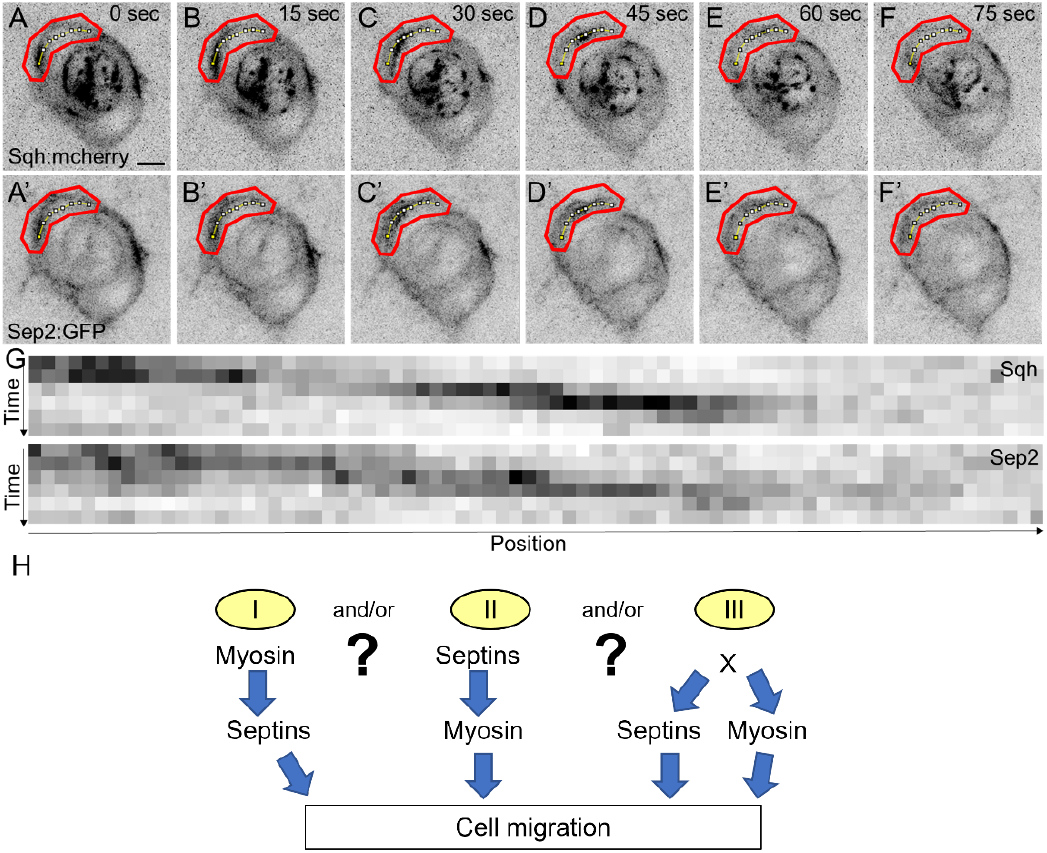
Septins and myosin localize dynamically. (A-F’) Max intensity stills of border cell migration with 15 second intervals between images labeled with Sqh:mcherry (A-F) and Sep2:GFP (A’-F’). (G) Kymographs of Sqh and Sep2, where the measured area is denoted by a dotted line, outlined in red in A-F’. (H) Three possible models to describe the relationship between septins and myosin in border cells. Scale bar in A is 5μm.

Three different mechanisms might account for septin and myosin colocalization (Figure 4H). By analogy to the bud neck in budding yeast (Lippincott and Li, 1998; Schneider et al., 2013), septin might recruit myosin. Alternatively, myosin might recruit septin. A third option is that both septins and myosin might both be recruited by an unknown upstream regulator (Figure 4H).

### Rho regulates septins and myosin independently

To test if myosin regulates septin expression or localization, we generated clones of cells expressing RNAi against Sqh in the follicular epithelium (Figures 5A-5B’) and in border cells (Figures 5C-5D’). We did not detect any change in Pnut expression level or localization in the basal follicular epithelium (Figure 5E) or in border cells (Figure 5F). Sqh activity is regulated by phosphorylation. To test whether Sqh activity state might affect septins, we expressed either a phosphomimetic (active) form of Sqh or a non-phosphorylatable (inactive) form. Neither had any discernible effect on Pnut expression or localization (Figures S5A-S5C). To test for epistasis between septins and myosin, we assessed whether myosin knockdown affected the Sep1 overexpression phenotype, and we found no change (Figures S5D-S5E’). This result suggests that the effect of septin on cluster shape is independent of myosin activity.

**Figure 5:**
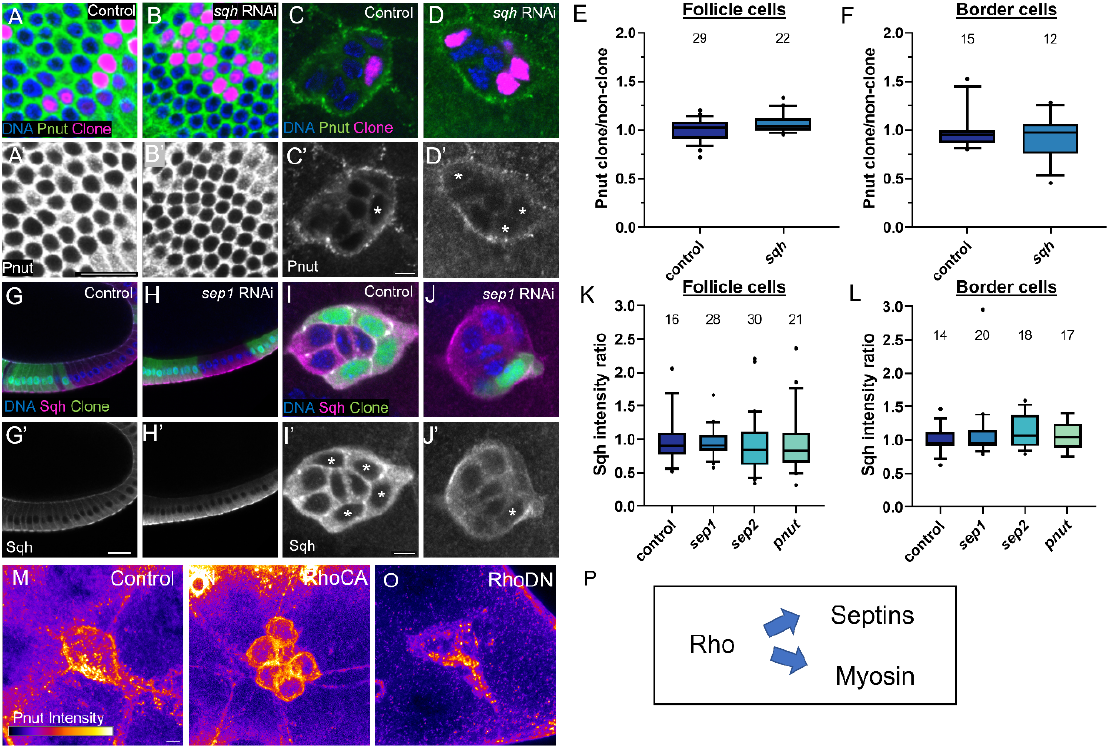
Rho regulates septins and myosin independently. (A-B) Single z-slice confocal images of epithelial follicle cells on the basal surface of the egg chamber clonally expressing UAS-sqhRNAi labeled with Hoechst (blue) and Pnut (green). Clones are marked with magenta nuclei, expressing nuclear-localized UAS-RedStinger. (A’-B’) Images from A-B for Pnut (gray). (C-D) Same as A-B but in border cells. (C’-D’) Same images from C-D but only labeled with Pnut (gray). Clones are marked with asterisks. (E-F) Box plots of Pnut intensity in clones compared to non-clones in follicle cells (E) and border cells (F). (G-H) Single slices of epithelial follicle cells clonally expressing UAS-septin RNAi and UAS-GFP labeled with Hoechst (blue). All cells express sqh:mcherry. (G’-H’) Same images from G-H but only labeled with Sqh (gray). (I-J) Same as G-H but in border cells. (I’-J’) Same images from I-J labeled with Sqh (gray). Clones are marked with asterisks. (K-L) Box plots of Sqh intensity in clones compared to non-clones in follicle cells (K) and border cells (L). (M-O) Max intensity projections of a control border cell cluster (M) or clusters expressing constitutively active Rho (N) or dominant negative Rho (O) with Pnut protein expression shown by a fire LUT. 15/15 RhoN19-expressing clusters showed cytoplasmic Pnut localization. 20/20 RhoV14-expressing clusters showed membrane localization for each of two transgenic lines tested. (P) The model best supported by our data, suggesting that Rho is upstream of septins and myosin. Scale bars in A’ and G’ are 20μm and scale bars in C’, I’, and M are 5μm.

To test whether septin regulates myosin, we clonally knocked down septin in the follicle cells (Figures 5G-5H’) and border cells (Figures 5I-5J’) in a Sqh:mCherry background and detected no difference in fluorescence intensity (Figure 5K and 5L). Furthermore, we could detect no time delay between the appearance of septin and myosin flashes. Together, these data suggest that septins and myosin might be recruited simultaneously by a common upstream regulator.

The GTPase Rho stood out as a potential candidate, as it is a well-established regulator of myosin activity. Rho has also been linked to septins in rat embryonic fibroblast cells (Ito et al., 2005) and animal cell cytokinesis (Carim et al., 2020). To test the effect of Rho activity on septin, we expressed constitutively active (V14) or dominant negative (N19) versions of Rho in the border cell cluster and stained for Pnut. Pnut typically accumulates at the border cell cortex and in a cytoplasmic pool (Figure 5M). In 100% of RhoV14-expressing border cells, we observed a marked increase in cortical accumulation of Pnut, at the expense of the cytoplasmic pool (Figure 5N). RhoV14 causes each individual border cell to round up, so the effect on septin localization could in principle have been an indirect effect of the change in cell shape. However, dominant negative Rho caused the opposite effect: septin relocalized from the cortex into the cytoplasm in 100% of clusters examined (Figure 5O). We conclude that active Rho recruits Pnut to the plasma membrane (Figure 5P), and thus septins are a previously unknown downstream effector of Rho in border cell migration.

### Septin knockdown and overexpression alter cluster shape

Border cell clusters change shape dynamically as they move. To initiate migration, one or two cells extend and retract dynamic, forward-directed protrusions (Fulga and Rørth, 2002; Murphy and Montell, 1996; Prasad and Montell, 2007). When a single leader is established, it communicates direction to the rest of the cells (Cai et al., 2014), inhibiting them from forming leader-like protrusions. Followers instead engage in crawling behaviors that both promote cluster cohesion and movement (Campanale et al., 2022). Leader protrusions act as sensory structures, probing the environment for chemical and physical cues (Dai et al., 2020; Mishra et al., 2019b). We noticed that clusters with reduced septin expression adopted abnormal shapes, particularly ectopic protrusions (Figure 6B), similar to the phenotype previously described for inhibition of myosin heavy or light chain (Mishra et al., 2019b). Conversely, overexpression of septin resulted in a rounder morphology (Figure 6C).

**Figure 6:**
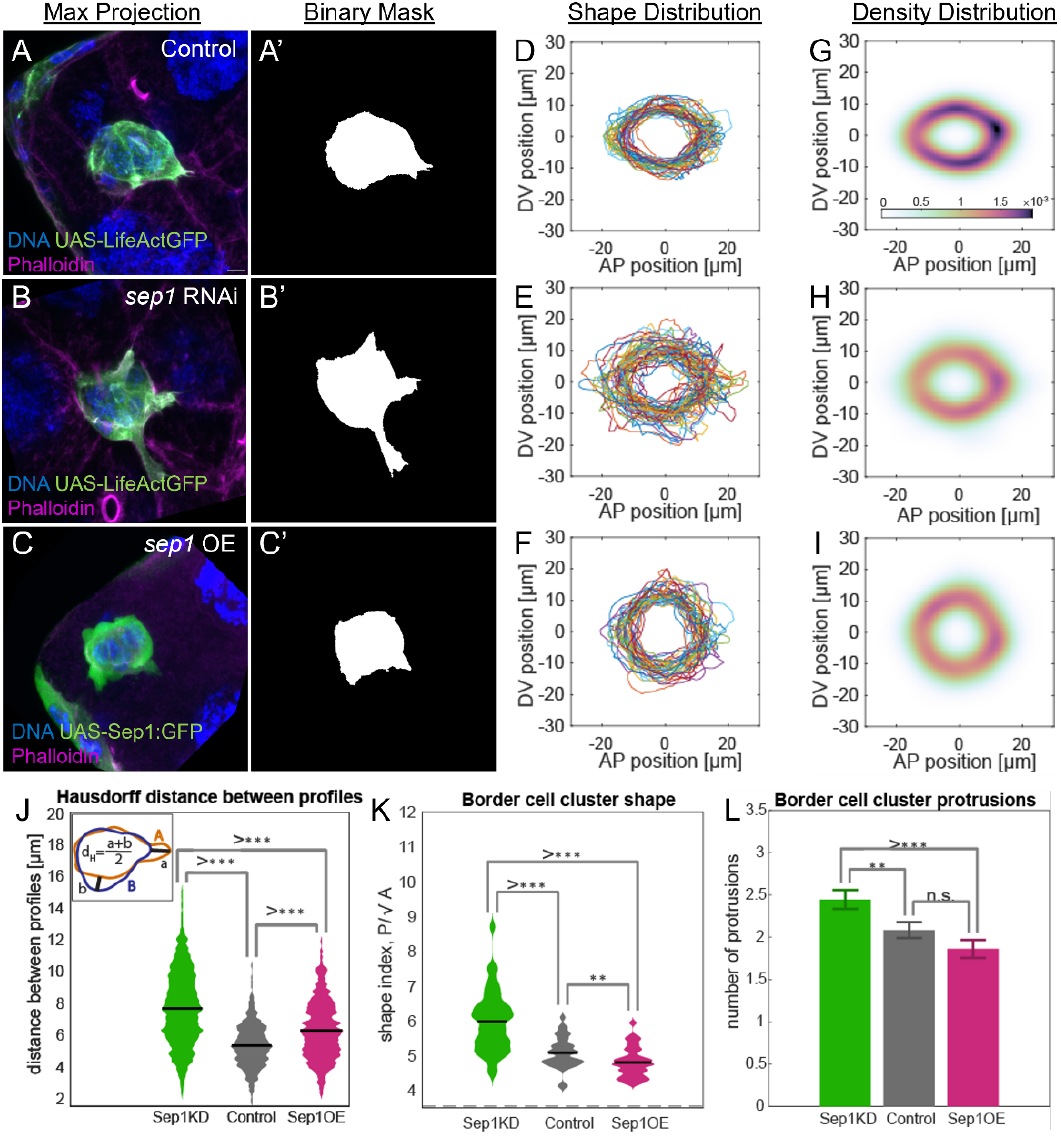
Septin knockdown and overexpression alter cluster shape. (A-C) Max intensity projections of border cell clusters expressing UAS-whiteRNAi (control) (A), septin1RNAi (B), or septin 1 overexpression (C). (A-B) Images labeled with UAS-LifeActGFP, Hoechst (blue), and Phalloidin (magenta). (C) Image labeled with UAS-septin1:GFP, Hoechst (blue), and Phalloidin (magenta). (A’-C’) Binary masks of each cluster in A-C. (D-F) Overlapping outlines of all clusters analyzed. D is control, E is septin knockdown, and F is septin overexpression. Axes are of anterior-posterior (AP) and dorsal-ventral (DV) position. (G-I) Density distribution plots of each genotype from (A-C) using a Gaussian Kernel Density Estimation. Density correlates with the color scale. (J) Dot plot of the Hausdorff distance between each genotype. Inset shows the calculation of Hausdorff distance as (a+b)/2. (K) Dot plot of the border cell cluster shape of each genotype. Shape index represents the perimeter divided by the square root of the area for each cluster. (L) Bar graph of the protrusion number for each genotype. (J-L) ** represent P<0.01 and *** represent P<0.001 when analyzed by a Kolmogorov–Smirnov test (J and K) or a one-sided t-test (L). Middle bars show means (J and K). Error bars show standard error (L). n=36 for control, 45 for septin knockdown, and 36 for septin overexpression.

To gain deeper insight into the normal functions of septins in controlling cluster shape, we compared cluster geometries as well as the shape variation within and between control, knockdown, and overexpression groups. We first converted maximum intensity projections of border cell clusters (Figures 6A-6C) to binary masks to define the cluster boundaries (Figures 6A’-6C’ and Figures S6A-S6O). We then overlaid the shapes to reveal the distributions in control (Figure 6D), septin knockdown (Figure 6E) and septin overexpression (Figure 6F). Control clusters had similar shapes (Figure 6D), whereas septin knockdown cluster shapes were more variable (Figure 6E). Compared to the control shape distribution, this distribution is more chaotic, with protrusions appearing in all orientations (Figures S6A-S6O). Septin overexpressing clusters also show more variability compared to controls (Figure 6F).

Another way to illustrate this variability is to display probability density distributions, which show that control clusters (Figure 6G) exhibit a more stereotyped set of shapes compared to the septin knockdown (Figure 6H). Clusters overexpressing septin likewise show more diffuse density distributions (Figure 6I). These clusters are rounder and less protrusive than controls, but some were skewed vertically and some horizontally relative to the direction of migration, increasing the overall variability and decreasing the density. To quantify the variability in cluster shapes, we calculated the Hausdorff distances as a measure of how far apart different curves (in this case the cluster perimeters) are from one another (Figure 6J and see Supplementary Information). Septin knockdown or overexpression results in larger and more variable Hausdorff distances compared to the control (Figure 6J), confirming that the shapes are more variable.

To quantify differences in shape, we measured the perimeter and divided it by the square root of the area to calculate a commonly used shape index (Bi et al., 2015). While a perfect circle minimizes the shape index at value of ∼3.5 (dashed line at bottom of Figure 6K), protrusions or irregular surfaces will register higher values. Septin knockdown clusters deviated the most from a circle while septin overexpressing clusters were significantly rounder than controls (Figure 6K). We then defined an objective method for counting protrusions based on the amount of extension from a reference circle (see Supplementary Information). Septin knockdown caused significantly more protrusions than control clusters, which in turn produced more protrusions than clusters overexpressing septin (Figure 6L). We conclude that clusters with altered septin expression show a greater variability in shape compared to control clusters, but knockdown causes excess protrusions while overexpression produces rounder clusters.

### Septin knockdown and overexpression alter surface texture

When Septin protein is added to the outside of giant unilamellar lipid vesicles *in vitro*, it is sufficient to deform the sphere and cause projections (Beber et al., 2019), so septins can act directly on membranes to reshape them. Septins are also known to induce membrane curvature in cells (Woods and Gladfelter, 2021). To describe membrane curvature in border cell clusters and determine if septin knockdown or overexpression altered curvature, we took advantage of a MATLAB toolbox for tissue cartography called ImSAnE (Image Surface Analysis Environment) (Heemskerk and Streichan, 2015). This allowed us to generate 3D models of the cluster surface from high-resolution Airyscan images. We noticed that septin knockdown clusters exhibited an overall smoother texture (Figures 7A-7A’) compared to control clusters (Figures 7B-7B’), with fewer sharp features, whereas clusters overexpressing septin had a rougher texture with more numerous and smaller domains of steep, positive and negative curvature (Figures 7C-7C’).

**Figure 7:**
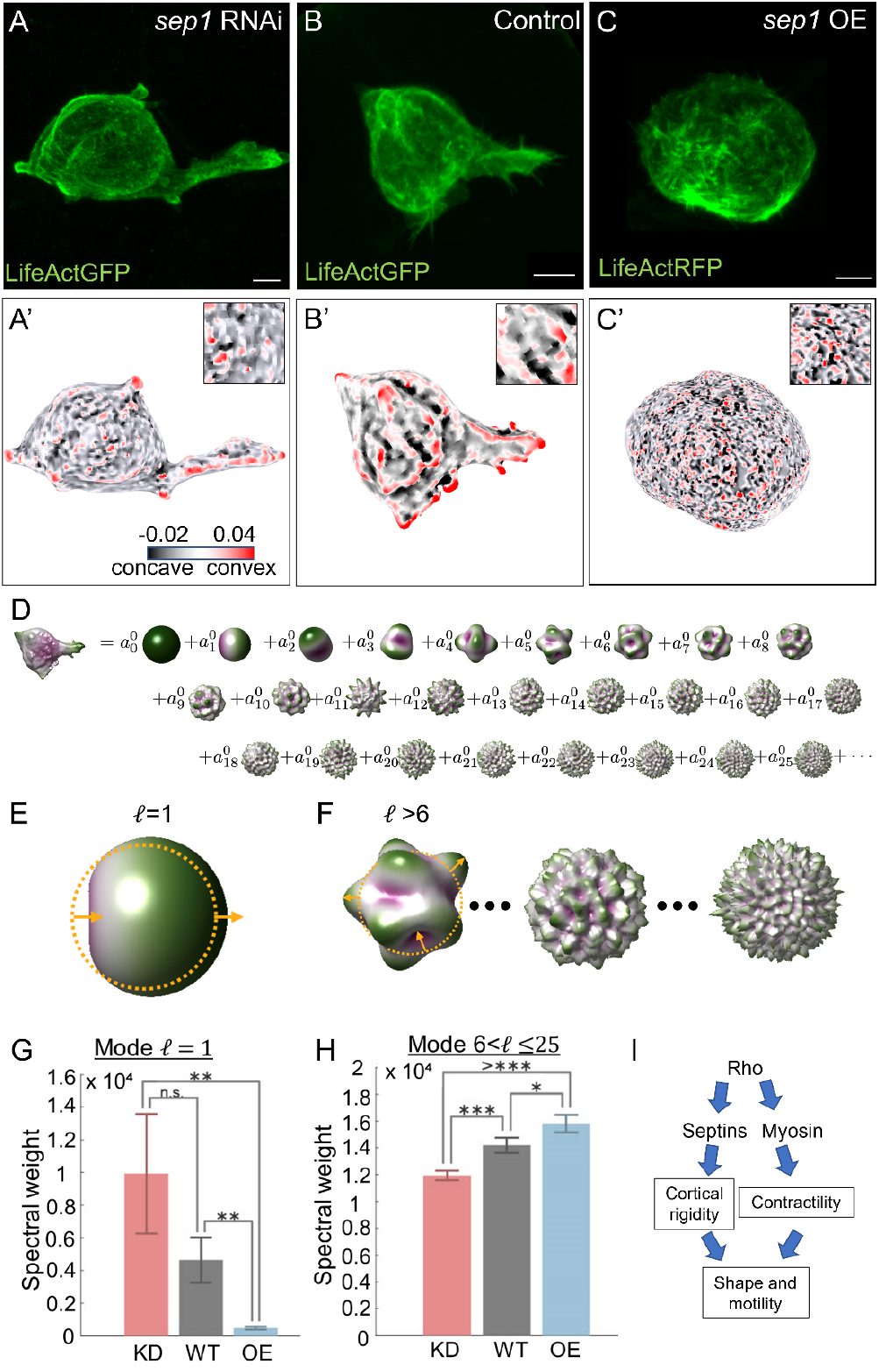
Septin knockdown and overexpression alter surface geometry. (A-C) Max intensity projection images of border cell clusters imaged by Airyscan after septin knockdown (A), control (B), or septin overexpression (C) and labeled by UAS-LifeActGFP (A-B) or UAS-LifeActRFP (C). (A’-C’) 3D curvature models of the surfaces of border cell clusters shown in A-C generated by a tissue cartography tool, ImSAnE (Image Surface Analysis Environment). Insets are zooms. Concave and convex surfaces are denoted by a black to red color scheme. (D-F) Spectral decomposition of 3D surface shape demonstrates that protrusivity decreases with septin expression, while surface roughness increases with septin expression. (D) The border cell cluster is a summation of component shapes of different weights. (E) The ℓ=1 spectral component measures ‘protrusivity’, or how much one side of the surface protrudes relative to the other with respect to a centroid found by mean curvature flow (see SI for details). (F) Higher-degree spectral components (sketched for three surfaces ℓ>6) measure finer-scale surface roughness. (G) Septin expression is inversely correlated with protrusivity. (H) Septin expression is correlated with greater surface roughness. (G-H) Error bars represent standard error. n=4 for control, 10 for knockdown, and 3 for overexpression. (I) Model of septins and myosin independently regulated by Rho.

To quantify these differences, we performed a spectral decomposition of the shapes using a sphere as a frame of reference (Dalmasso et al., 2022; Mitchell and Cislo, 2022) (see Supplementary Information). This analysis describes a complex 3D object such as a border cell cluster as a sum of contributions of component shapes (Figure 7D), which are each weighted in the sum. The first component (which carries a weight A1) measures deviations from sphericity at the largest possible scale – the cluster diameter. This component therefore describes an asymmetric shape with a single protrusion (with respect to a suitably-defined centroid as described in the Supplementary Information) (Figure 7D). Higher-degree spectral components measure the contributions of shapes with finer-scale surface roughness (Figures 7D-7F). We found that the septin knockdown clusters, and to a lesser degree controls, are best described by heavily weighting the contribution of the first component and thus have a higher spectral weight in the first degree (Figure 7G), whereas the shapes of clusters overexpressing septin are best described by heavily weighting the higher degree components (shown in Figure 7H for weights A7-25). With increasing septin expression, the higher-degree components contribute more to cluster shapes (Figure S7A-S7B), demonstrating that higher septin expression results in a fine-scale roughness.

Together the analyses of cluster geometry suggest that the fine scale smoothness caused by septin knockdown results in more variability in cluster shape, more large-scale protrusions, and an overall more asymmetric shape. Conversely, septin overexpression causes fine scale roughness that results in an overall more spherical and symmetrical cluster shape. While smoothness might naively be taken as an indication of rigidity, these findings suggest the membrane is more rigid when corrugated by septin expression, akin to how fine-scale undulations in cardboard prevent bending at larger scales (see discussion).

## Discussion

Here we show that active Rho recruits septins to the plasma membrane where the level of septin tunes surface properties, independently from myosin. Both myosin-mediated contractility and septin-mediated rigidity ensure appropriate protrusion and promote collective migration through the confined 3D environment of the egg chamber (Figure 7I), addressing key open questions in the septin field.

### Septin function in border cell migration

Border cell migration is a well-studied example of *in vivo* collective cell migration where the roles of Rho GTPases (Colombié et al., 2017; Murphy and Montell, 1996) and myosin (Aranjuez et al., 2016; Chen et al., 2020; Combedazou et al., 2017; Majumder et al., 2012; Mishra et al., 2019b; Zeledon et al., 2019) have been extensively explored. To date, myosin has been the only Rho effector identified in these cells. Here we show that active Rho recruits septins to the plasma membrane where they co-localize dynamically with myosin. Septins interact directly with lipids, and can deform membranes of Giant Unilamellar Vesicles and induce spikes (Beber et al., 2019; Tanaka-Takiguchi et al., 2009). Septins can also dramatically remodel 2D lipid bilayers into 3D structures (Vial et al., 2021). Septins are therefore most likely to exert their effects on border cells at membranes, so membrane recruitment is a key step in regulating septin function.

We also show that either increasing or decreasing the level of septin expression alters cluster shape and impairs motility. We propose that the concentration of septin protein affects the deformability of the cell surface: low septin expression increases protrusions whereas increasing septin expression decreases protrusions. Interestingly, this effect on large scale protrusions likely results from the effects of septins on fine scale roughness vs smoothness of the cell surface in a somewhat surprising way. The rough texture of the surfaces of clusters over-expressing septins may function like the corrugations in reinforced cardboard where fine scale roughness makes the material more rigid and inhibits larger scale deformation. Thus septin overexpression causes the overall cluster shape to be more spherical, whereas the fine-scale smoothness of septin knockdown clusters renders the surface more deformable and thus excessively protrusive.

Septins are required in a great diversity of cell types for many different behaviors. It is striking that the morphology of border cells lacking septin exhibits commonalities with septin knockdown phenotypes in a variety of individually migrating cells including the distal tip cell and neuronal axons in C. elegans (Finger et al., 2003) and mammalian T-cells (Tooley et al., 2009). In these examples, as in border cells, septin knockdown caused extra and ectopic protrusions as well as impaired migration. However, these studies did not report the consequences of septin overexpression.

Tooley et. al, found that T cells with reduced septin expression are better able to squeeze into confined spaces that are smaller than the diameter of the cell even though septin-depleted cells were worse at persistent and directional migration. Border cell clusters squeeze into spaces much smaller than even a single border cell, so one might expect that reduced septin would facilitate their movement. However, they also have to migrate directionally and persistently. Hence, perhaps, the need for precisely optimal levels of septin expression.

Neuronal progenitors in the developing mammalian cerebral cortex migrate along radial glial fibers by extending a long, leading process. Septin 14 or Septin 4 knockdown causes the neuronal processes to be shorter and impairs migration (Shinoda et al., 2010). Exactly what role septins play in this context is not yet clear. Furthermore, the effects of septin overexpression have not been analyzed in these contexts, but based on our findings, we would predict that overexpression would cause different changes to cell shapes and would also likely impair migration.

Our analysis shows that the precise level of septin expression can affect the domains of microcurvature in cells with increasing septin leading to less deformable membranes. Overexpression and loss-of-function of septins have been associated with a great variety of human diseases (Marttinen et al., 2015; Montagna et al., 2015; Peterson and Petty, 2010; Wang et al., 2018), though precisely how changes in septin expression cause disease remain largely mysterious. Here we show how reduction or increase in septin expression produces opposite effects on cell surface properties and cell shape and that both impair motility. Humans have a larger set of septin genes and proteins than *Drosophila*. By modulating not only the expression levels but the compositions of septin filaments, this larger set might allow for even greater diversity of membrane properties and thus more diverse and complex cell shapes and behaviors.

## Materials and Methods

### Drosophila genetics

Fly strains used in this study are listed in Supplementary Table 1. Detailed fly genotypes in each experiment are listed in Supplementary Table 2.

### Fly Husbandry

Fly strains were raised in vials containing a standard cornmeal-yeast food (https://bdsc.indiana.edu/information/recipes/molassesfood.html) which contains 163g yellow cornmeal, 33g dried yeast, 200mL molasses and 16g agar with 2.66L water. All flies were raised in vials containing 5mL fly food.

### RNAi knockdown with Gal4 drivers

2-4 day-old females were kept in 29C for 3 days, transferred to a vial with dry yeast each day until dissection. FLPout clones were first heat-shocked for one hour at 37C to induce clones, kept at room temperature for 8 hours, heat-shocked again at 37C for one hour, and then kept at 29C for 3 days with dry yeast until dissection.

### Egg chamber dissection and staining

Adult female ovaries were dissected in Schneider’s Drosophila medium (Thermo Fisher Scientific, Waltham, MA; 21720) with 20% fetal bovine serum. Ovarioles containing egg chambers of the desired stages were pulled out of the muscle sheath with #55 forceps.

For fixed sample staining, ovarioles were then fixed for 15 min in 4% paraformaldehyde. After fixation, ovarioles were washed with PBS/0.1% Triton X-100 (PBST) or PBS/0.4% Triton X-100 (PBST), and then incubated with primary antibodies overnight at 4 °C. The following day, ovarioles were washed with PBST before incubation in secondary antibodies and Hoechst overnight at 4 °C. The following day, ovarioles were again washed with PBST. Samples were stored in VECTASHIELD (Vector Laboratories, Burlingame, CA) at 4 °C before mounting.

The following antibodies and dyes were used in this study: Hoechst (1:1000, sigma-aldrich), rat anti-E-cadherin (1:25, DCAD2, DSHB), mouse anti-Pnut (1:50, 4C9H4, DSHB), rabbit anti-GFP (1:300, lifetech), rabbit anti-mCherry (1:500, novusbio), Alexa 488, 568, 647 (1:200, lifetech), phalloidin 647 (1:200, sigma-aldrich).

### Fixed Sample Imaging and Image Processing

Samples were mounted on a glass slide in VECTASHIELD. Images were taken on a Zeiss LSM 800 confocal microscope, using a 20×1.2 N.A. objective, 40×1.4 N.A. water objective, or 63x, 0.8 NA oil objective. images were taken on a Zeiss LSM 800 confocal microscope, using 63x, 0.8NA oil objective.

### Live Imaging

Ovaries were dissected in Schneider’s Drosophila medium (Thermo Fisher Scientific, Waltham, MA) with 20% fetal bovine serum. Individual ovarioles were carefully pulled out and stage 9 egg chambers were removed. The egg chambers were collected in a 0.6 mL tube and washed with dissecting medium twice, then added to 100 uL dissecting medium with insulin (100 ug/uL) and 1% low melt agarose. 100 uL medium with the egg chambers then were mounted on a 50mmLumox dish. Time-lapse imaging was performed using a 20×1.2N.A. objective or 40×1.1 NA water immersion objective lens.

### Tissue Cartography and Curvature 3D Models

We imaged migrating border cell clusters through high-resolution airyscan imaging. These images were imported into Ilastik (1), an opensource software for segmenting cells using machine learning. We used this to define the surface of the border cell cluster, and exported this file as a .h5 file. This .h5 file was imported into meshlab (2) to clean up the mesh as well as generate a file that can be analyzed using ImSAnE (Image Surface Analysis Environment) (6). Meshlab version used was MeshLab_64bit v1.3.3. Mesh construction was done by: 1) importing the cell surface; 2) Filters -> Sampling -> Poisson disc sampling. Base mesh subsampling option must be checked. Number of samples used was 15,000; 3) Filters - > Normals, Curvature and Orientation -> computing 7ormal for point sets; 4) Filters -> Remeshing, Simplification and Reconstruction -> surface reconstruction: poisson. The reconstructed surface is then exported as a PLY file with the flags and 7ormal data included.

The PLY file was analyzed using ImSAnE, details provided in the reference above as well as comments within the example scripts provided by the authors in their github [https://github.com/idse/imsane]. Specifically we modified the example script TutorialIlastikDetectorSpherelikeFitter.m running on Matlab_R2019a.

## QUANTIFICATIONS AND STATISTICAL ANALYSES

### Migration Defect Quantification

For quantification of migration defects, stage 10 egg chambers were scored by eye. The position of the border cell clusters were assigned to categories of 0%, 1-25%, 26-50%, 51-75%, or 76-100% based on their distance from the anterior of the egg chamber to the oocyte.

### Colocalization Quantifications

For quantification of Sep2 and Pnut and LifeActGFP controls, high-resolution Airyscan z-stack images of the border cell cluster were taken at 63x magnification with the 0.8 NA oil objective. A single slice in the center of the z-stack was selected. A ROI was drawn to outline the cluster and background was subtracted. Colocalization of two channels within the ROI was measured using FIJI Coloc2 function (FIJI function: Analyze > Colocalization Analysis > Coloc 2). Pearson’s correlation, r was used as the readout of colozalization.

### Pnut intensity Quantifications

For quantification of Pnut intensity, 20x, 40x, or 63x images of Pnut channel were measured in FIJI. When measuring Pnut intensity in follicle cells or border cells, a single slice in the center of the z stack was measured that clearly showed the cells of interest. About 3-5 follicle cells or one border cell were measured when comparing clonal cells to adjacent non-clones (Figure 3A-K). All quantifications use raw integrated density normalized to area.

### Sep2:GFP intensity Quantifications

For quantification of Sep2:GFP intensity, samples were probed with anti-GFP antibody. 20x images of GFP channel were measured in FIJI. When measuring Sep2:GFP intensity in follicle cells, a single slice in the center of the z stack was measured that clearly showed the cells of interest. bout 3-5 follicle cells were measured when comparing UAS-RNAi expressing cells to control cells. All quantifications use raw integrated density normalized to area. All quantifications use integrated density.

### Geomstats Analysis in Python

We plotted the shapes of each genotype using a software package in Python called Geomstats to project each cluster shape onto the manifold of discrete curves. On the manifold of discrete curves, each point represents a different cluster shape. Thus, when we “projected” a cluster to the manifold, we are identifying which point on the manifold matches the shape of that cluster. We oriented all clusters so that the direction of motion was facing right. Shape indices were calculated by measuring the perimeter of each cluster (segmented as a maximum intensity projection) and dividing it by the square root of the area of that shape. To count protrusions, we identified stable maxima that are at least a distance R away from the centroid of the segmented border cell cluster and have a peak that is at least 0.3*R larger than the surrounding data to avoid overcounting. Here, we set R to equal the radius of a circle whose area is equal to the cluster under consideration. To measure Hausdorff distance between curves, we first collected all aligned curves outlining the border cell clusters, resampled them at 100 approximately equally spaced points, and expressed the positions of these points in physical units relative to the center of the cluster. For each pair of curves A and B within a given genotype, we then found the Hausdorff distance d(A,B) which is the maximum distance of any point in curve A from its closest point in B. The measurement was then symmetrized: 0.5*(d(A,B) + d(B, A)) for statistical analysis because the measurement of d(A,B) is not independent of d(B,A). We performed an all-to-all comparison within each category (WT, KD, OE) to build up a distribution. This gave N*(N-1)/2 symmetrized distances between pairs of curves.

### Spectral Power Analysis

To quantify the change in surface geometry in septin knockdowns and overexpression, we decomposed each cell cluster’s surface profile into spherical harmonics using methods published in (Mitchell and Cislo, 2022).

This measures the amount of surface deformation as a function of spatial scale, separating slow, long-wavelength variations in protrusion/ingression from short-wavelength roughness. This procedure is analogous to taking a Fourier transform, in which a signal is decomposed into components which vary in their wavelength. To facilitate comparison across cells and across conditions, we conformally mapped each cell to a sphere (Kazhdan et al., 2012) and defined a field on this sphere which indicates the extent of radial protrusion or ingression.

In detail, after acquiring each cell surface as a triangulated mesh, we conformally mapped a given cell’s mesh to a spherical surface via mean curvature flow according to Kazhdan et. al, resulting in a sphere of radius ***R*** centered at ***r***_**0**_. Let us d Let us denote this conformal map enote this conformal map ***f***. This mapping enabled comparison across samples and across genotypes. We found the original distance of all surface vertices from the center ***r***_**0**_ (defined via conformal mean curvature flow) and subtracted ***R***, which is the mapped radial distance from ***r***_**0**_. This gave us a measure of how much the surface is extended beyond ***R*** or retracted from ***R*** at each point on the mesh. This defined a ‘radial’ surface profile ***δr***, which can be expressed either as a function ***δr*(*x*)** of position on the cell surface, or as a function ***δr*(*f*(*x*)))** of position on the mapped sphere found by mean curvature flow. Scalar field patterns on the sphere can be compared directly across samples. Therefore, we decomposed the scalar field defined on the mapped (spherical) surface into spherical harmonics. Spherical harmonics 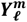 are a set of functions that can reproduce the original field when multiplied by appropriate spectral weights 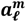 and summed together:

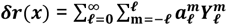

More formally, they are eigenmodes of the Laplace-Beltrami operator defined on the sphere. The inner product between a given eigenmode and the measured radial surface profile field ***δr*** defines the spectral weight at that spatial scale. We then plotted these weights as a function of index ℓ. Weights of different spherical harmonics 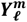 with identical index ℓ are summed together in each bin, since they represent similar wavelengths of variation along the surface:

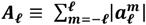

We took the absolute value of the weights in the sum. We then plotted ***A***_***ℓ***_ as a function of the degree ℓ.

Comparing the lowest anisotropic component ℓ = 1 across samples showed a significant trend, with spectral weight anti-correlated with septin expression. This component has higher weight when one side of the cell protrudes relative to the other. Higher-degree components measure short-wavelength roughness of the surface, and most high order modes are individually correlated with septin expression. Summing many modes together resulted in significant correlation with septin expression, and this result is insensitive to the lower or upper bound on ℓ included in the sum.

### Statistics and Data Presentation

Standard statistical tests were performed using Graphpad Prism and MATLAB. Ordinary one-way ANOVA, followed by Tukey’s multiple comparisons test was used for comparing multiple groups with similar variance as determined by Brown–Forsythe test for Figure 3. Kolmogorov– Smirnov test was used in Figures 6J and 6L. One-sided t-test was used in Figures 6K, 7F, and 7G. One asterisk denotes a significance of P<0.05, two asterisks represent P<0.01, and three asterisks are P<0.001. Graphs in Figures 1, 2, 3, and 5 were generated using Graphpad Prism. Plots in Figure 6 were generated through MATLAB and Python. Plots in Figure 7 were made using MATLAB and Graphic. All confocal images belonging to the same experiment were acquired using the exact same settings. For visualization purposes, brightness adjustments were applied using FIJI to the confocal images shown in the figure panels. All quantitative analyses were carried out on unadjusted raw images. All fly crosses were repeated at least twice and ovary dissections and staining were repeated at least three times. Sample size was not predetermined by statistical methods but we used prior knowledge to estimate minimum sample size. The experiments were not randomized. Investigators were not blinded.

## End Matter

### Author Contributions and Notes

Experiments were designed by A.M. Gabbert., J.A. Mondo, J.P. Campanale, and D.J. Montell. Experiments were carried out by A.M. Gabbert. J.A. Mondo and J.P. Campanale assisted with computer software and microscopy. Data analysis was performed by A.M. Gabbert. Geomstats analysis was performed by N. Mitchell, A. Myers, and N. Miolane. Spectral power analysis was performed by N. Mitchell. This manuscript was prepared by A.M. Gabbert., J.A. Mondo, J.P. Campanale, N. Mitchell, and D.J. Montell.

The authors declare no conflict of interest.

All data are available in the manuscript or the supplementary materials. Materials available upon request.

## Acknowledgments

We thank the entire Montell lab for discussions and feedback. We thank Miles Keats, Nick Keefer, Abraham Sontay, and Spencer Phillips for technical assistance and Dillon Cislo for discussions on spectral analysis. Funding: This work was supported by NIH grant GM073164 to D.J.M, ACS grant PF-17-024-01-CSM to J.P.C, and support from the Helen Hay Whitney Foundation to N.P.M. This work was also supported in part by the National Science Foundation Grant No. NSF PHY-1748958 to the Kavli Institute for Theoretical Physics. We thank the Developmental Studies Hybridoma Bank for providing antibodies and the Bloomington Drosophila Stock Center and the Vienna Drosophila Resource Center for providing fly stocks.

---

**Supplementary Figure 1.**
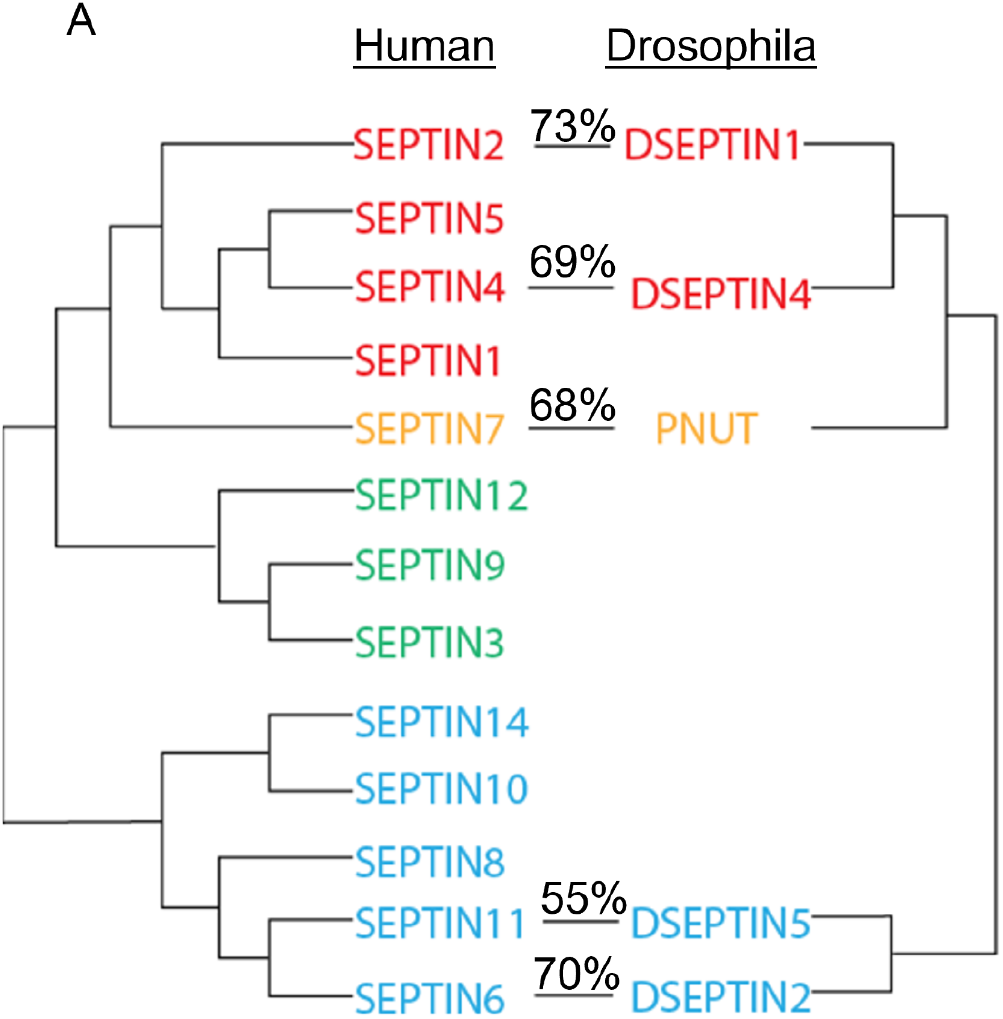
*Drosophila* septins have high similarity to their human orthologs. (A) Humans have 13 septins while *Drosophila* have 5. Calculated values denote the similarity between amino acid sequences for human and Drosophila septin homologs.

**Supplementary Figure 2.**
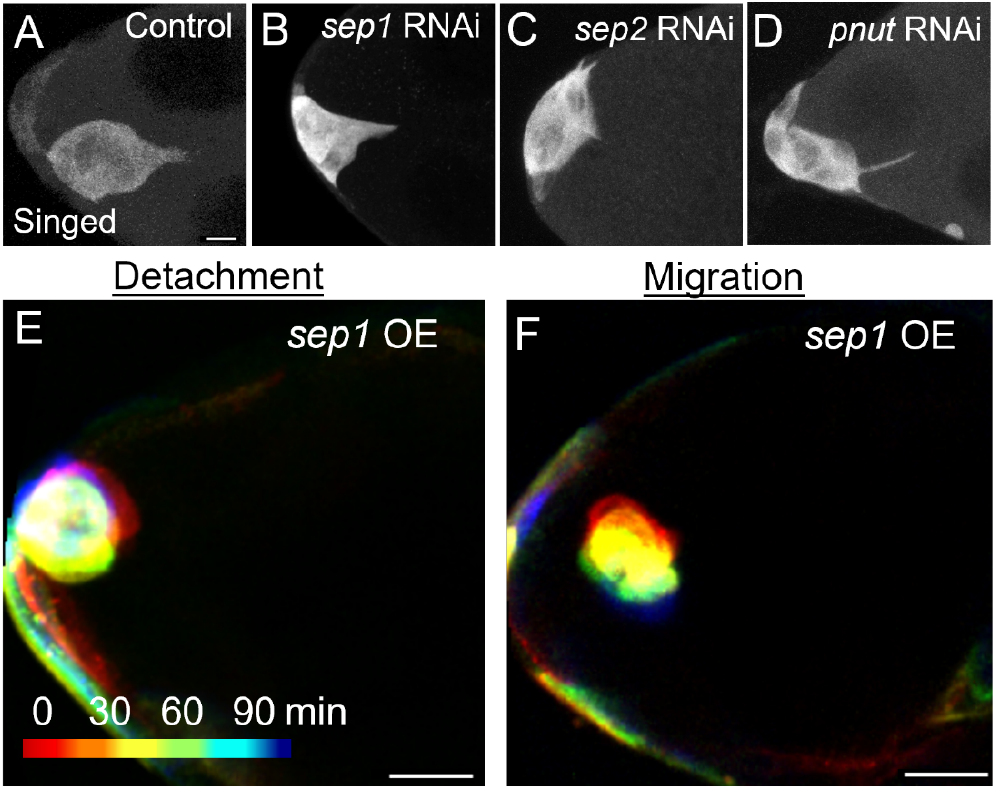
(A-D) Max intensity projections of border cell clusters expressing UAS-whiteRNAi (control) or septinRNAi with c306Gal4, labeled with Singed (gray). (E-F) Max intensity projections of four time points of detachment (E) or migration (F) from timelapse series of UAS-Sep1:GFP (overexpression) clusters. The scale bar in A, E, and F represent 20μm.

**Supplementary Figure 3.**
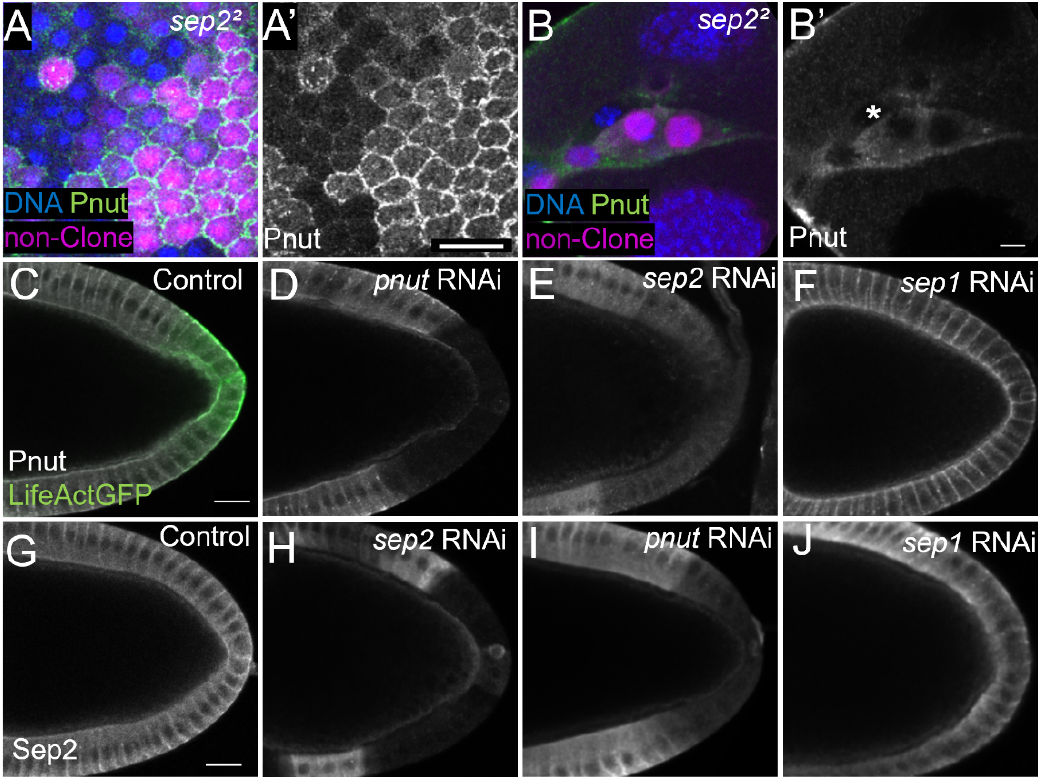
(A-B) Single z-slice images of epithelial follicle cells or border cells clonally expressing a null mutation for Septin 2 labeled with Hoechst (blue) and Pnut (green). Negative clones are marked with magenta nuclei (RFP), representing wildtype cells. Cells lacking magenta nuclei lack Septin 2, expressing a null mutation.. (A’-B’) Same images from A-B but only labeled with Pnut (gray). A Sep2 mutant border cell clone is marked with an asterisk. (C-J) Single z-slice images of epithelial follicle cells in the center plane of the egg chamber expressing septin RNAi in the posterior follicle cells, shown by UAS-LifeActGFP in C. All cells are stained with Pnut (gray) (C-F) or express Sep2:GFP (gray) (G-J). The scale bars in A’, C, and G are 20μm. The scale bar in B’ is 5μm.

**Supplementary Figure 4.**
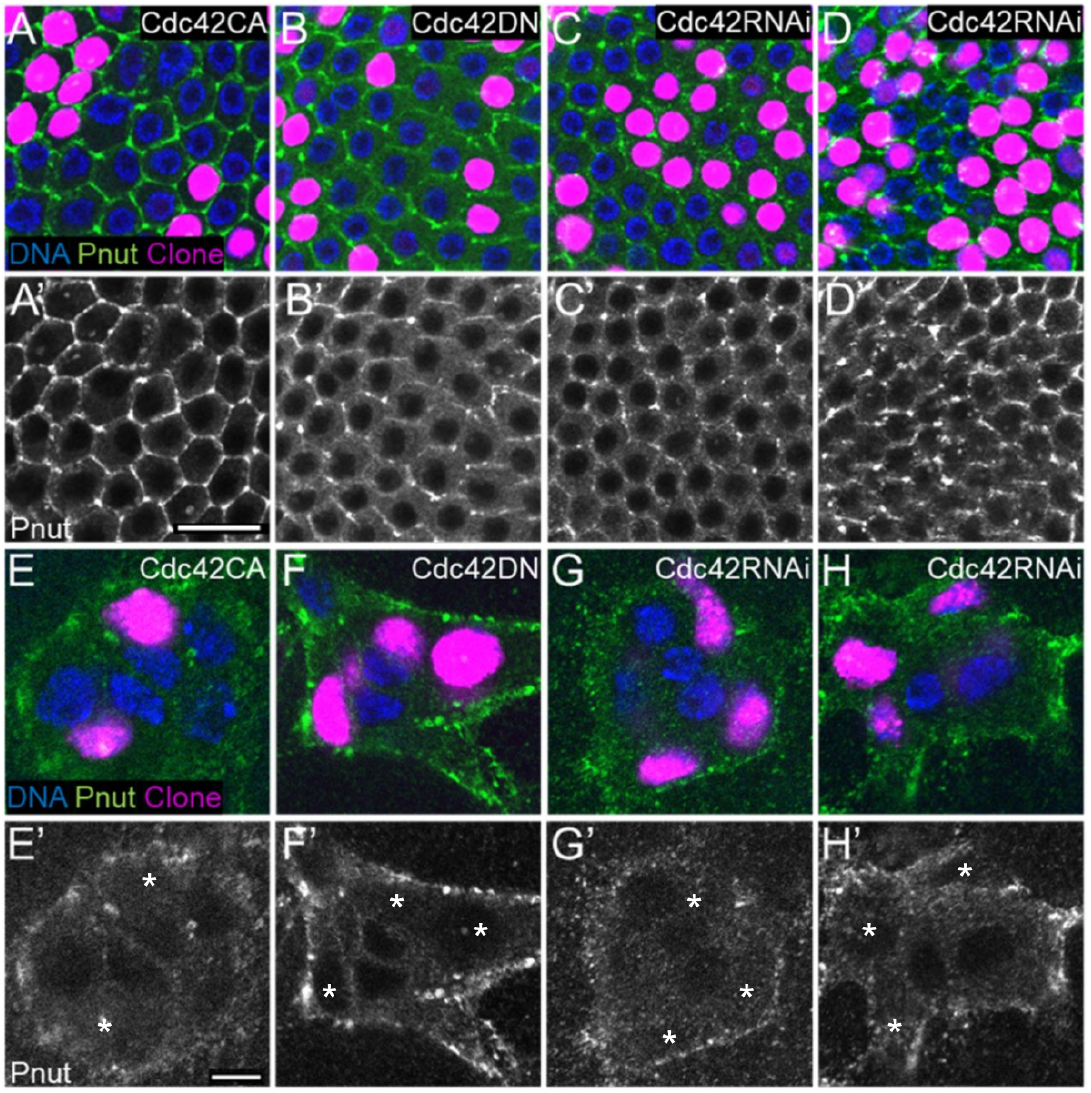
(A-D) Single slice images of epithelial follicle cells on the basal surface of the egg chamber clonally expressing constitutively active Cdc42 (A), a dominant-negative version of Cdc42 (B), or one of two Cdc42 RNAi constructs (C-D) labeled with Hoechst (blue) and Pnut (green). Clones are marked with magenta nuclei, expressing nuclear-localized RedStinger. (A’-D’) Same images from A-D but only labeled with Pnut (gray). (E-H) Same as A-D but in border cells. Clones are marked with asterisks. (E’-H’) Same images from E-H but only labeled with Pnut (gray).The scale bar in A’ is 20μm and the scale bar in E’ is 5μm.

**Supplementary Figure 5:**
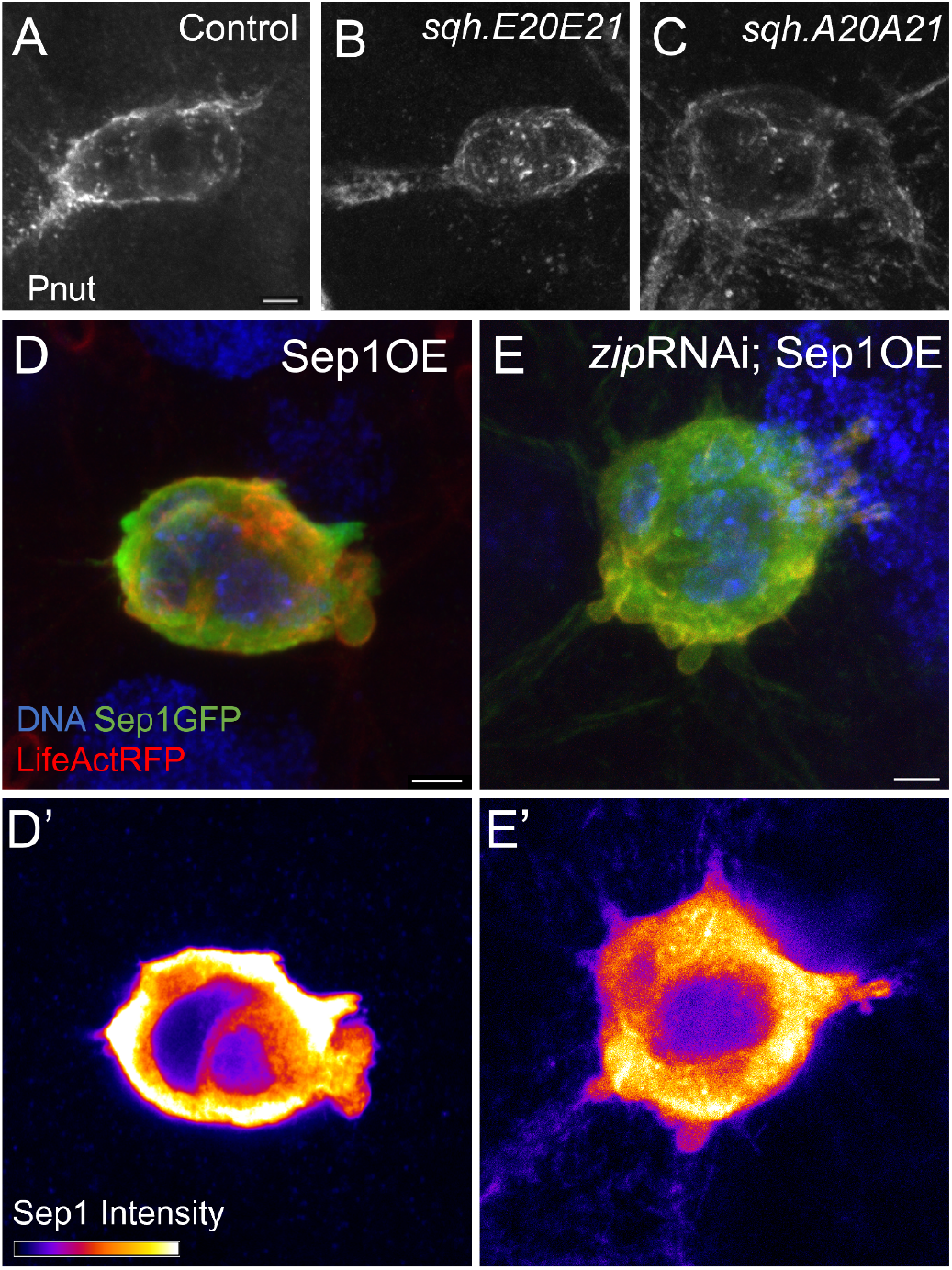
(A-C) Max intensity projections of border cell clusters expressing UAS-whiteRNAi (control) (A), UAS-sqhE20E21 (B), or UAS-sqhA20A21 (C) labeled with Pnut (gray). (D-E) Max intensity projections of border cell clusters expressing Sep1:GFP and UAS-LifeActRFP, or the addition of UAS-zipRNAi (E) labeled with Hoechst (blue). (D’-E’) Single slice images of (D-E) with Sep1:GFP shown by a fire color gradient. The scale bars in A, D, and E are 5μm.

**Supplementary Figure 6.**
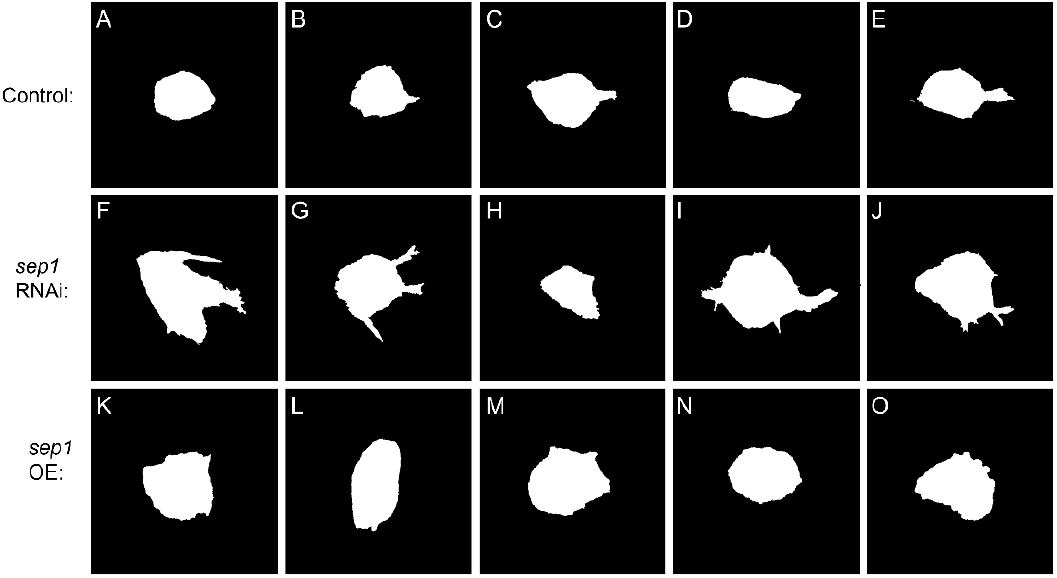
(A-O) Black and white binary masks of border cell clusters, showing additional examples of control (A-E), Septin 1 knockdown (F-J) and Septin 1 overexpression (K-O).

**Supplementary Figure 7.**
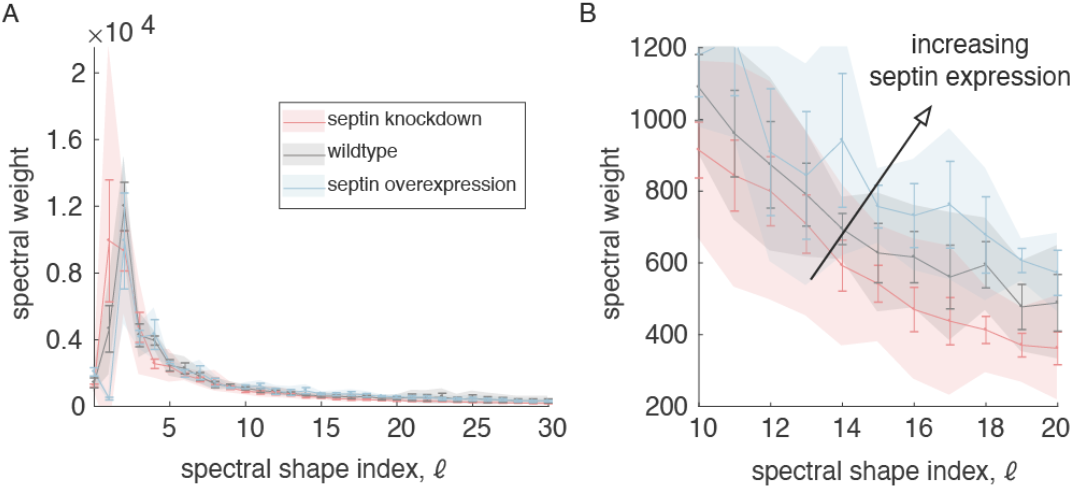
Decomposing cell surface profiles into spherical harmonics yields a spectral weight associated with discrete spatial scales. Qualitatively, this measures the contribution of each spatial scale in the overall shape of the cell, where higher degrees ℓ – correspond to finer scale roughness of the surface. (A) We plot Aℓ m= -ℓℓ |aℓm| as a function of degree ℓ, where aℓm is the spectral weight associated with each spherical harmonic Yℓm. Bars represent standard error and the shading represents standard deviation. (B) Zooming in on spectral weights for higher-degree components shows correlation of spectral weight with septin expression.

**Supplementary Table 1.**

| Genotype | Source | Identifier |
| --- | --- | --- |
| c306Gal4;UAS-LifeActGFP;Gal80ts | Denise Montell Lab Stock, University of California Santa Barbara | N/A |
| y[1] w[*]; P{w[+mC]=Septin2-GFP.SG}3 | Bloomington Drosophila Stock Center | Cat#BDSC_26257 |
| hsFLP;AyGal4-25b,UASredstingerNLS | Denise Montell Lab Stock, University of California Santa Barbara | N/A |
| c306Gal4;sqh::sqh-mcherry/CyO | Denise Montell Lab Stock, University of California Santa Barbara | N/A |
| sqh::sqh-mcherry/(CyO);MKRS/TM6 B | Denise Montell Lab Stock, University of California Santa Barbara | N/A |
| D. melanogaster y[1] v[1]; P{y[+t7.7] v[+t1.8]=TRiP.HMS00017}at tP2 | Bloomington Drosophila Stock Center | Cat#BDSC_33623 |
| w[1118]; P{GD8198}v17344 | Vienna Drosophila Resource Center | Cat#VDRC_17344 |
| w1118; P{GD11240}v26413 | Vienna Drosophila Resource Center | Cat#VDRC_26413 |
| w1118 P{GD1512}v11791 | Vienna Drosophila Resource Center | Cat#VDRC 11791 |
| w[1118]; P{w[+mC]=UAS-Cdc42.V12}LL1 | Bloomington Drosophila Stock Center | Cat#BDSC_4854 |
| w[*]; P{w[+mC]=UAS-Cdc42.N17}3 | Bloomington Drosophila Stock Center | Cat#BDSC_6288 |
| P{VSH330192}attP40 | Vienna Drosophila Resource Center | Cat#VDRC_330192 |
| P{KK108698}VIE-260B | Vienna Drosophila Resource Center | Cat#VDRC_100794 |
| w[*]; P{w[+mC]=UASp-Septin1.GFP}3 | Bloomington Drosophila Stock Center | Cat#BDSC_51346 |
| w[*]; P{w[+mC]=UASp-Septin2.O}18A/CyO | Bloomington Drosophila Stock Center | Cat#BDSC_91012 |
| w1118;<br>P{GD1695}v7917/TM3 | Vienna Drosophila Resource Center | Cat#VDRC_7917 |
| w[*]; P{w[+mC]=UAS-Rho1.V14}5.1/CyO | Bloomington Drosophila Stock Center | Cat#BDSC_7330 |
| w[*]; P{w[+mC]=UAS-Rho1.V14}2.1/TM6 | Bloomington Drosophila Stock Center | Cat#BDSC_8144 |
| P{w[+mC]=UAS-Rho1.N19}1.3, w[*] | Bloomington Drosophila Stock Center | Cat#BDSC_#7327 |
| hsFLP;AyGal4, UAS-GFP;MKRS | Denise Montell Lab Stock, University of California Santa Barbara | N/A |
| w[*]; Septin2[1]/TM6B, Tb[1] | Bloomington Drosophila Stock Center | Cat#BDSC_91002 |
| w[*]; Septin2[2]/TM6B, Tb[1] | Bloomington Drosophila Stock Center | Cat#BDSC_91003 |
| yw, hsFLP;;FRT82B, ubiRFPnls | Denise Montell Lab Stock, University of California Santa Barbara | N/A |
| w[*]; P{w[+mC]=UASp-sqh.A20A21}3 | Bloomington Drosophila Stock Center | Cat#BDSC_64114 |
| w[*]; P{w[+mC]=UASp-sqh.E20E21}3 | Bloomington Drosophila Stock Center | Cat#BDSC_64411 |
| P{VSH330299}attP40 | Vienna Drosophila Resource Center | Cat#VDRC_330299 |

**Supplementary Table 2.**
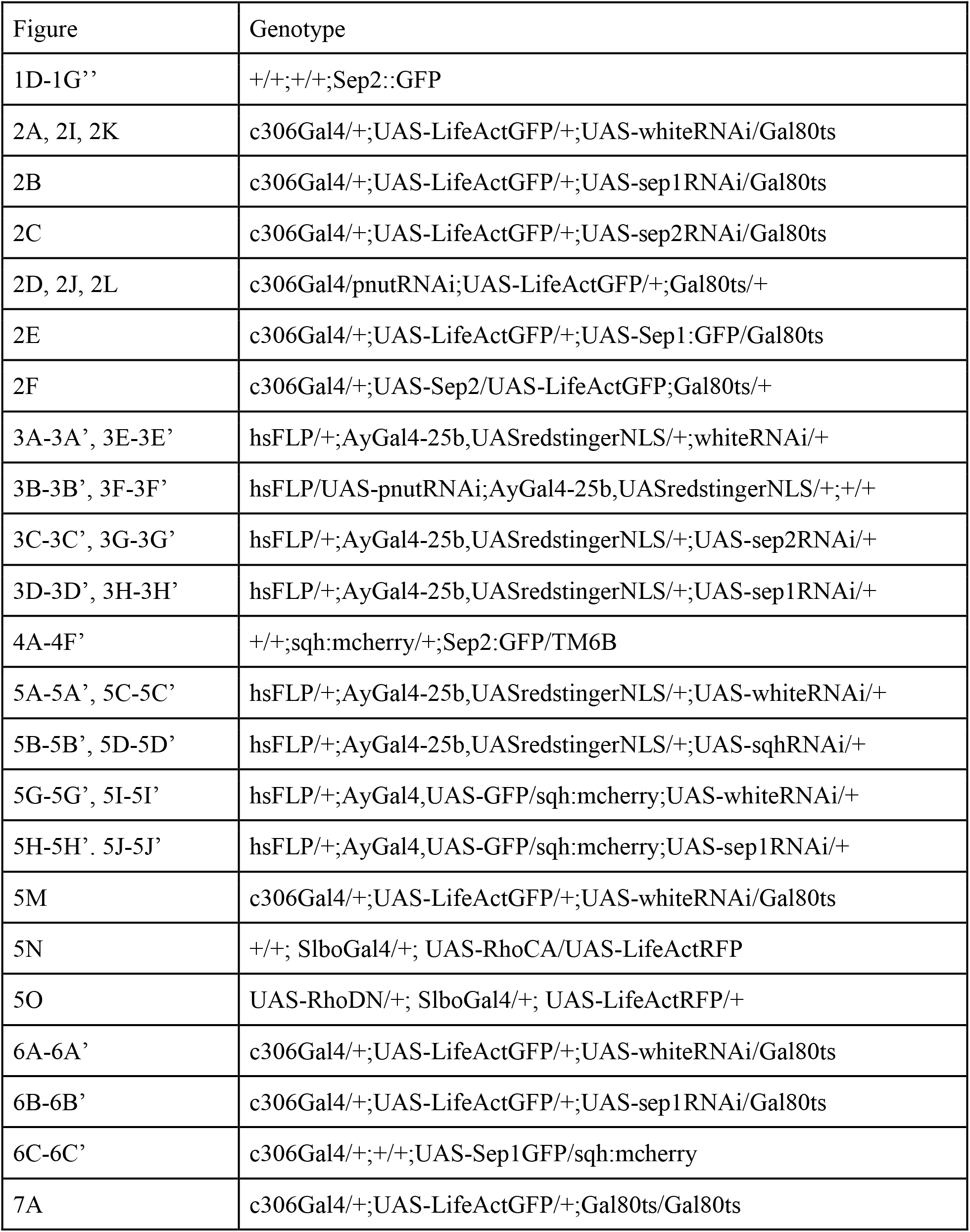
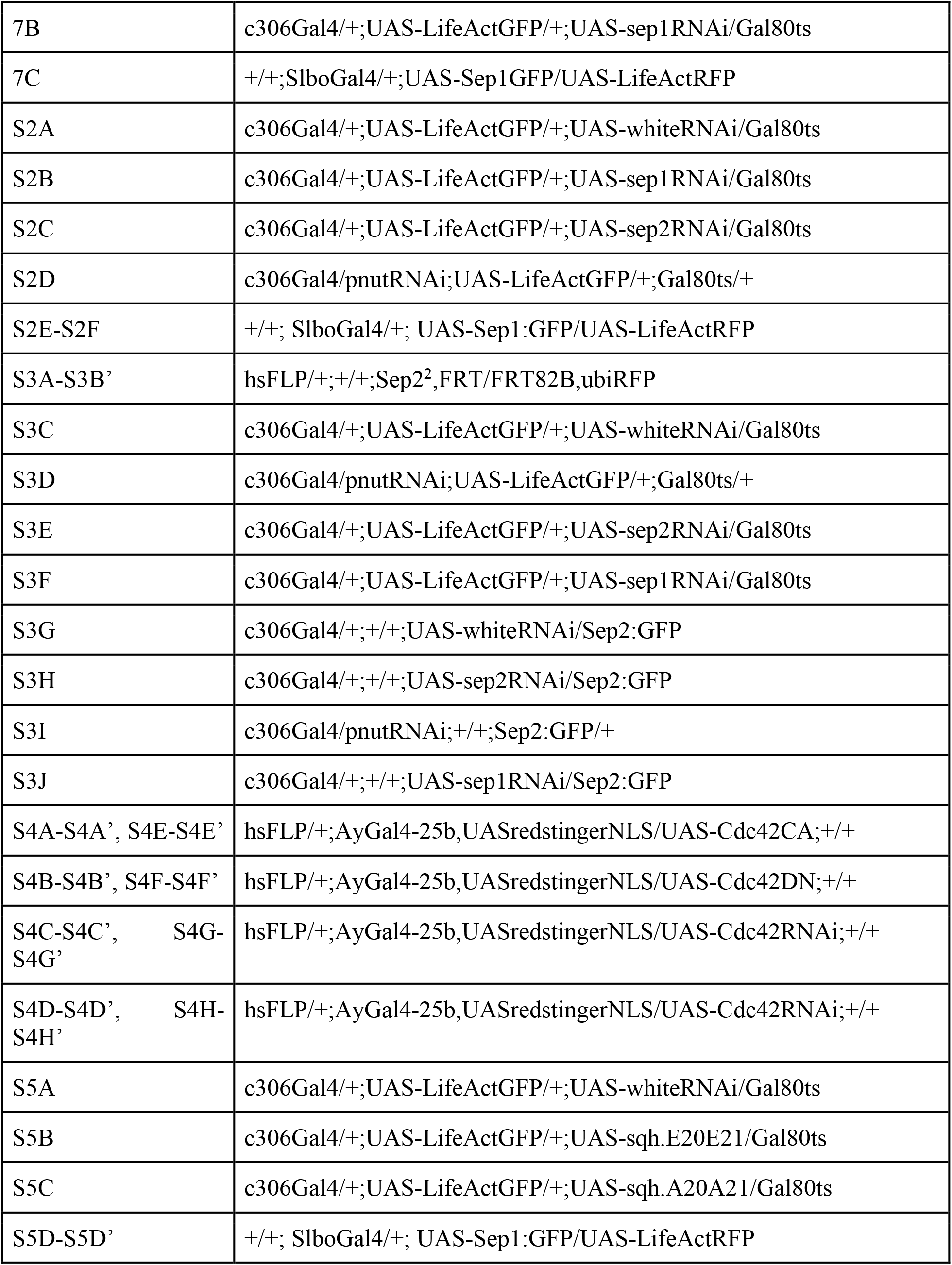
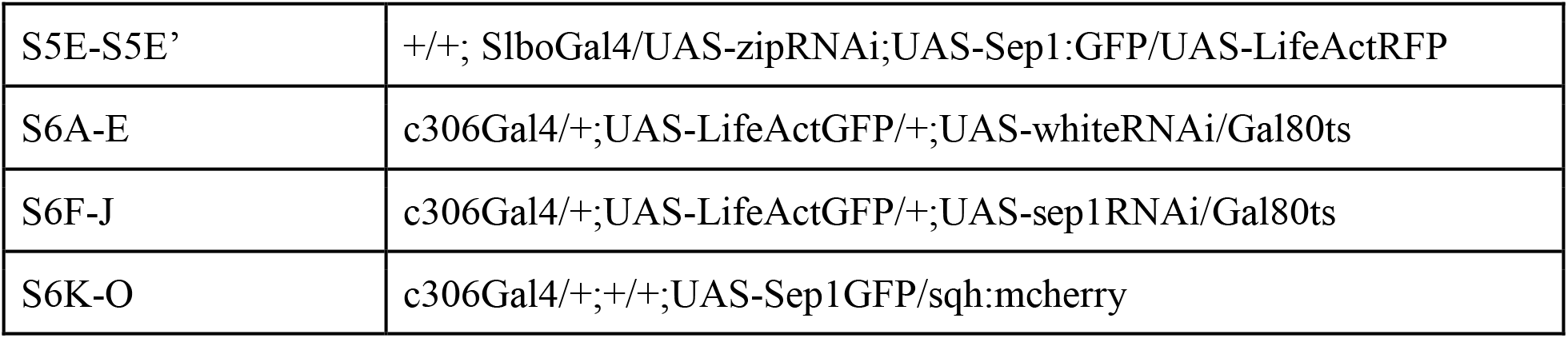

## Notes

### Competing Interest Statement

The authors have declared no competing interest.

### Summary of Updates

This version of the manuscript has been revised to include new results that show that septins are recruited to the border cell plasma membrane by active Rho, new formal mathematical analyses of the effects of septin knockdown and overexpression on small scale surface curvature and large scale protrusions. The results suggest that septin-induced micro-corrugations make the surface more rigid and difficult to deform at larger scales.

